# Asbestos accelerates disease onset in a genetic model of Malignant Pleural Mesothelioma

**DOI:** 10.1101/2020.06.16.154815

**Authors:** Pooyeh Farahmand, Katarina Gyuraszova, Claire Rooney, Ximena L. Raffo-Iraolagoitia, Emma Johnson, Tatiana Chernova, Tiziana Monteverde, Kevin Blyth, Roger Duffin, Leo M. Carlin, David Lewis, John le Quesne, Marion MacFarlane, Daniel J. Murphy

**Author notes:** Equal contribution (Pooyeh Farahmand, Katarina Gyuraszova). Present affiliation: Institute of Cancer Sciences, University of Glasgow (John le Quesne).

## Abstract

**Hypothesis:** Asbestos-driven inflammation contributes to malignant pleural mesothelioma beyond the acquisition of rate-limiting mutations.

**Methods:** Genetically modified conditional allelic mice that were previously shown to develop mesothelioma in the absence of exposure to asbestos were induced with lentiviral vector expressing Cre recombinase with and without intrapleural injection of amosite asbestos and monitored until symptoms required euthanasia. Resulting tumours were examined histologically and by immunohistochemistry for expression of lineage markers and immune cell infiltration.

**Results:** Injection of asbestos dramatically accelerated disease onset and end-stage tumour burden. Tumours initiated in the presence of asbestos showed increased macrophage infiltration. Single-agent suppression of macrophages in mice with established tumours failed to extend survival.

**Conclusion:** Asbestos-driven inflammation contributes to the severity of mesothelioma beyond the acquisition of rate-limiting mutations, however, targeted suppression of macrophages alone in established disease showed no therapeutic benefit.

## INTRODUCTION

Malignant pleural mesothelioma (MPM) is a devastating neoplasm arising in the lining of the chest cavity that is causally linked to asbestos inhalation (1). This cancer is characterised by a lengthy incubation period between exposure to asbestos and the onset of clinical symptoms, followed by very rapid progression from diagnosis to morbidity (2). Treatment options are severely limited and 5-year survival is <10% (3, 4). Although commercial use of asbestos is banned in most western countries, mining and use of asbestos continues unabated in several developing economies. In the UK and elsewhere, a legacy of asbestos-insulated buildings continues to drive new cases of MPM (5).

Direct inoculation of asbestos fibres into rodents has been used to model both early and late stage disease (6, 7). Injection of asbestos into the chest cavity provokes a rapid inflammatory response, comprised initially of neutrophils followed rapidly by macrophage infiltration (8). Fibre dimensions that thwart engulfment and clearance result in “frustrated phagocytosis” and the establishment of chronic inflammation (9). The resulting prolonged release of cytokines and reactive oxygen species are thought to create a permissive milieu that drives proliferation and mutation in pleural mesothelial cells (10, 11). Recent longitudinal analysis revealed steadily progressive disease in wild-type mice injected with long-fibre amosite asbestos, with epigenetic silencing of key tumour suppressor loci manifest in pre-neoplastic pleural lesions (7). Progression to mesothelioma however was reported in <10% of such mice after 18 months of incubation and the precise mechanism of progression to malignancy remains poorly understood.

The genetic landscape of end-stage human MPM is now well-defined. Inactivating mutations and deletions of 3 tumour suppressor loci, namely *CDKN2A/B, NF2* and *BAP1* predominate, with functional loss of each tumour suppressor occurring in over 50% of cases and single or combinatorial loss of the 3 loci found in all possible permutations (12, 13). Loss of additional tumour suppressors linked to the Hippo pathway (*LATS1, LATS2, MST1, SAV*), histone methylation (*SETD2, SETDB1*), RNA splicing (*SF3B1, DDX3X*) and mTORC1 regulation (*TSC1, TSC2, ULK2*) occur at lower frequencies. Inactivating mutations of *Tp53* are reported to be present in <10% of MPM but are associated with more aggressive disease (12). In contrast, activating mutations and amplification of known driver oncogenes common in other cancer are noticeably absent, although overexpression of hTERT, linked to mutation of the gene promoter region, has been reported (14).

Genetic deletion of *Tp53* (15), *Nf2* (16), *Cdkn2a* (17) and *Bap1* (18) have each been shown to predispose mice to asbestos-induced mesothelioma, albeit with incomplete penetrance and variable latencies. The use of constitutive (ie. whole organism) and in some instances heterozygous alleles likely masked the true susceptibility of tumour suppressor-deleted pleural mesothelium to carcinogenic transformation. The first purely genetic models of MPM demonstrated that combinatorial loss of tumour suppressors in the appropriate anatomic location sufficed to give rise to MPM in the absence of asbestos inoculation (19). Intrapleural delivery of Adeno-Cre was used to conditionally delete floxed alleles for *Nf2, Cdkn2a* and *Tp53* in various combinations, and gave rise predominantly to mesothelioma with sarcomatoid or biphasic histology. A somewhat confounding observation was the frequent development of tumours in unintended tissues: these included lymphomas, leiomyomas, tumours of unspecified origin and hepatomegaly. These off-target neoplasms suggest escape of the viral vector from the chest cavity with consequent Cre-mediated allele inactivation in the affected tissues. More recently, the same strategy was applied to mice bearing combinations of floxed alleles for *Nf2*, *Cdkn2a* and *Bap1* (20, 21). Although Cre-mediated deletion of any pair of homozygous alleles sufficed to give rise to pleural mesothelioma, acquired loss of Bap1 protein expression was found in the majority of *Nf2*^*FL*^;*Cdkn2a*^*FL*^ tumours *in vivo*, and deletion of all three was required for spheroid formation *in vitro* (20). As reported for the Tp53 study (19), tumour histology was predominantly sarcomatoid (20) or biphasic (21) and off-target tumours were again frequently detected in both studies.

Such *in vivo* studies unequivocally demonstrate a functional requirement for these tumour suppressors to prevent mesothelioma development. One question they fail to address however is whether asbestos-driven inflammation plays any role in MPM progression beyond its contribution to the acquisition of rate-limiting mutations. This question has immediate clinical relevance, as the persistence of asbestos fibres sustains an inflammatory microenvironment even after the acquisition of disease-limiting mutations and throughout therapeutic intervention. We therefore asked if intrapleural injection of disease-relevant doses of asbestos has any impact on disease progression in mice that are genetically destined to develop MPM. Our results show a dramatic acceleration of disease onset in such mice upon inclusion of asbestos exposure, with significant implications for pre-clinical modelling of this disease *in vivo*.

## RESULTS

### Asbestos exacerbates genetically driven Mesothelioma

To investigate the impact of asbestos on mesothelioma progression in mice that are genetically destined to develop mesothelioma, we used triple allelic mice previously described to be sufficient for mesothelioma development in the absence of fibre exposure: *Cdkn2a*^*−/−*^ mice bearing homozygous floxed alleles for *Nf2* and *Tp53* (*Cdkn2a*^*−/−*^;*Nf2*^*fl/fl*^;*Tp53*^*fl/fl*^ – abbreviated as CNP) (19). To initiate tumours in the appropriate anatomic location (ie. the pleural cavity), adult mice triple homozygous for all 3 alleles were injected intrapleurally with 10^7^ pfu of lentiviral vector expressing Cre recombinase. After a 10-day interval to minimise the potential for viral infection of infiltrating leukocytes, mice were similarly injected once with low dose (25μg) long fibre amosite asbestos or left untreated (Figure 1A). Mice were monitored for development of clinical symptoms and euthanised for tissue collection at pre-defined humane end-points. CNP mice induced with lenti-Cre all developed clinical symptoms, notably including hiccups and laboured breathing. Strikingly, mice that were also administered asbestos exhibited dramatically accelerated disease, requiring euthanasia at a median of 88 days post Cre-mediated induction, as compared with those that did not receive asbestos, which had a median survival of 148 days post induction (Figure 1B). Asbestos administration to uninduced CNP mice did not result in any mice developing symptoms of disease within the same timeframe, in agreement with previous reports of similarly low dose asbestos treatment of wild-type mice (7). Upon dissection, frank tumours were clearly visible on the mesothelial surfaces of the lungs, heart, diaphragm and rib cages of all lenti-Cre induced mice, consistent with presentation in human mesothelioma patients. Tumour burden was visibly more pronounced in mice that also received asbestos compared with those that did not (Figure 1C). Tumours were detectable by MRI and moderately glucose avid and could be detected at all sites in end-stage mice by [^18^F]-FDG-PET and autoradiography (Figure S1). Notably, off-target tumours in other tissues, such as those reported in similar allelic mice induced using high doses of adenoviral-Cre vectors (19–21), were not detected, either in the presence or absence of asbestos treatment.

**Figure 1.**
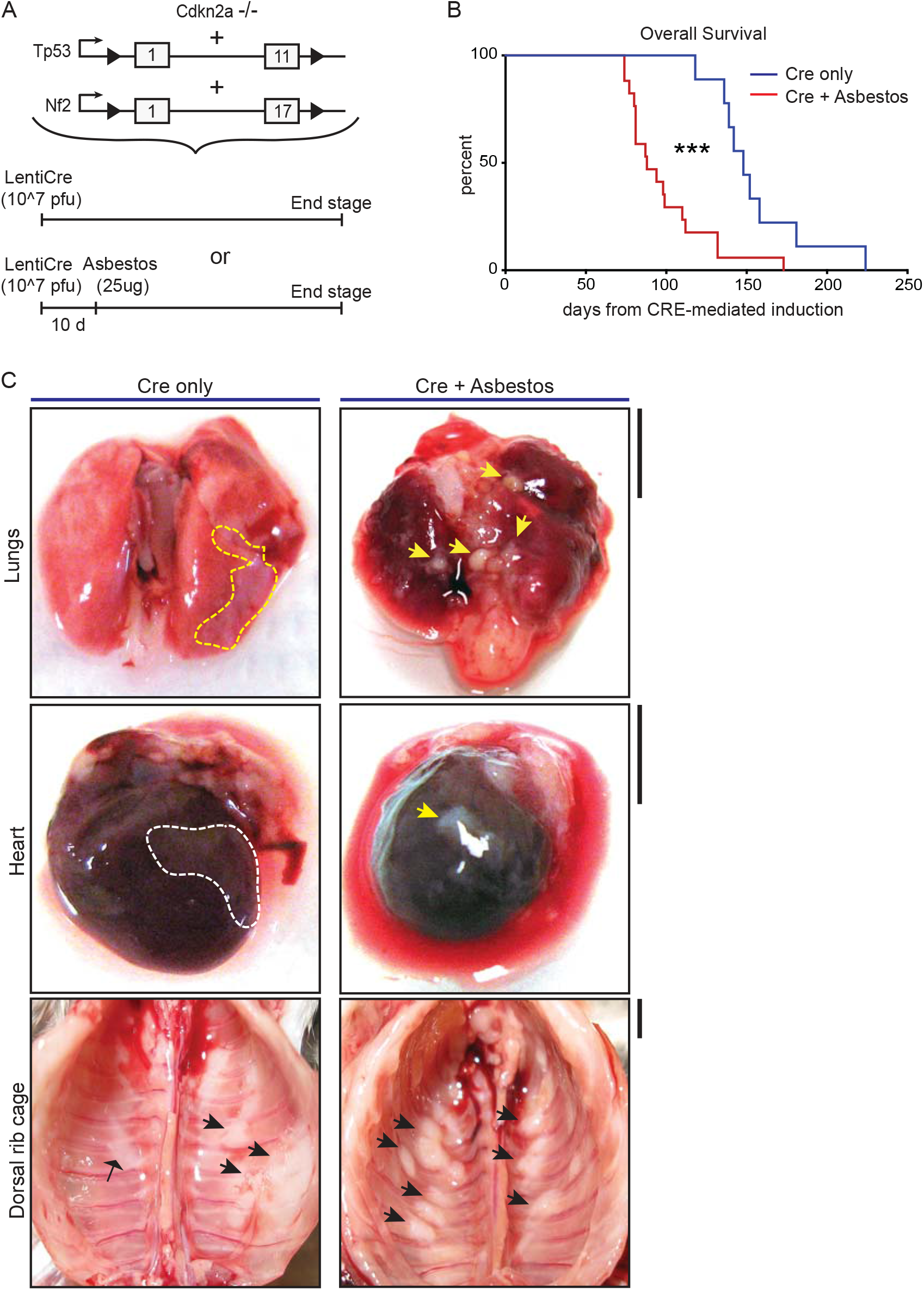
Asbestos injection reduces survival of CNP mice. **A)** Schematic of the alleles and induction strategies employed in this study. **B)** Kaplan Meier plot of overall survival of lenti-Cre induced CNP mice with (N=17) or without (N=9) asbestos injection. *** denotes p<0.001 (Mantel Cox logrank test). **C)** Representative photographs showing anatomic distribution of visible MPM lesions (arrowheads) in mice from (B). Scale bars = 4mm.

### Histological characterization of lenti-Cre and/or asbestos induced lesions

Harvested organs presenting with visible disease were sectioned and stained with H&E for histological examination. The visceral and parietal pleura of uninduced CNP mice injected with asbestos alone and harvested 10 months later all showed discrete regions of reactive thickening, reflective of chronic inflammatory disease, but no tumour formation. In contrast, histological examination confirmed the presence of malignancy in all CNP mice induced with lenti-Cre. Tumours arising in mice induced with Cre alone tended to grow laterally and often contained cystic or luminal structures, whereas those induced by the combination of Cre + asbestos typically formed densely packed spheroids, often with a necrotic core. Marked invasion into cardiac muscle and diaphragm was evident in both tumour-prone cohorts but was more commonly observed in mice that received the combination of Cre + asbestos (Figure 2A). Human MPM is a heterogenous disease with three main histological subtypes, depending on the predominant cellular component and biological behaviour. The epithelioid subtype accounts for 50-70% of cases, 10-20% of cases are sarcomatoid, and biphasic accounts for approximately 30% of reported cases (22). Tumours arising in CNP mice were predominantly (>90%) epithelioid with a small subset of lesions showing discrete regions of sarcomatoid morphology (Figure 2B).

**Figure 2.**
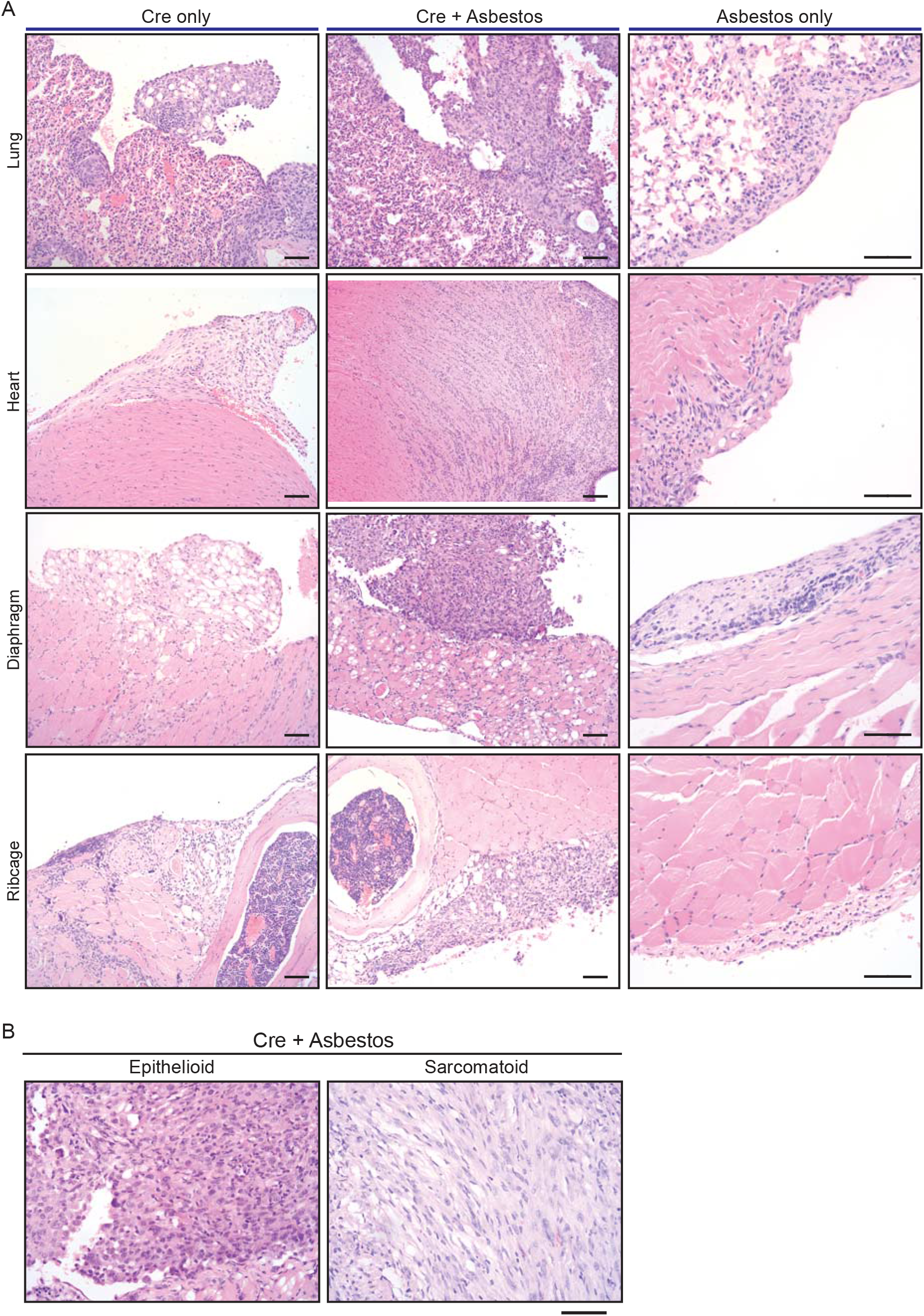
Histology of mesothelial lesions in CNP mice induced with lenti-Cre and/or asbestos. **A)** Representative images of indicated H&E stained tissues from mice induced with lenti-Cre (N=XX), Cre + asbestos (N=YY), both at end-stage, or asbestos without Cre-mediated allele induction, harvested 10 months after asbestos injection (N=3). Scale bars = 100μm. [check] **B)** H&E-stained examples of epithelioid and sarcomatoid histologies of tumours from CNP mice induced with lenti-Cre + asbestos. Scale bars = 100μm.

### Asbestos does not overtly alter the tumour-cell intrinsic mesothelioma phenotype

We asked if the inclusion of asbestos manifestly altered the tumour cell-autonomous characteristics of mesothelioma arising in our GE mouse model. For this analysis we focussed on tumours arising from the diaphragm mesothelium, given that decalcification protocols required for analysis of the chest wall can adversely impact immunohistochemical staining. It should be noted that tumours arising on the diaphragm were histologically indistinguishable from those arising on the lung or mediastinum. Tumours of approximately equal size arising in lenti-Cre induced mice with and without asbestos were compared by IHC for expression of cell lineage markers (pan-cytokeratin, vimentin and smooth muscle actin) along with Ki67 and the commonly used mesothelioma marker WT1. Individual tumours stained positively for all such markers to varying degrees, however averaging IHC marker expression across tumours in individual mice showed no consistent difference in expression of WT1 or Ki67, suggesting that the inclusion of asbestos does not overtly influence the tumour phenotype (Figure 3A & B).

**Figure 3.**
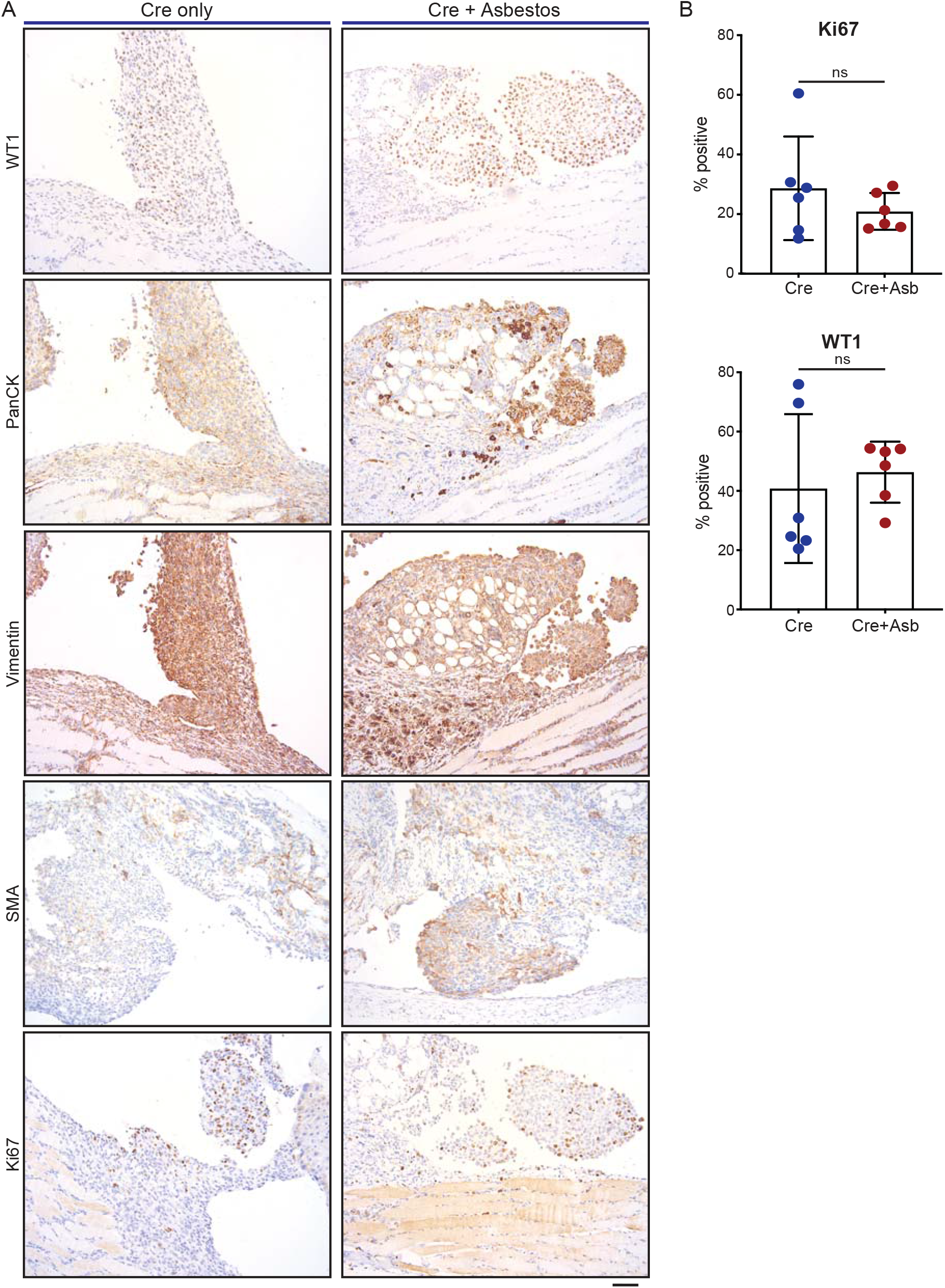
Asbestos does not alter the tumour phenotype of induced CNP mice. **A)** Representative IHC staining of tumours arising from diaphragm mesothelium in CNP mice induced by lenti-Cre, with and without asbestos. Scale bar = 100μm. **B)** Quantification of Ki67 (top panel) and WT1 (bottom panel) expression in diaphragm tumours as per (A). N=6 per cohort; ns = not statistically significant.

### Asbestos dramatically increases macrophage infiltration of tumours

Human pleural mesothelioma is characterised by a pronounced immunosuppressive inflammatory microenvironment (23–26). Recent work has demonstrated the presence of all major immune populations in both pre-malignant lesions of asbestos-exposed wild-type mice (7) and in end-stage tumours of GE mice without asbestos exposure (21). We therefore investigated the impact of asbestos on major tumour-infiltrating immune populations in our GE mouse model. End-stage tumours on the diaphragms of CNP mice induced with Cre +/− asbestos were stained for expression of CD8, CD4, CD45R (aka B220, expressed on B lymphocytes), NKp46, F4/80 (macrophages) and Ly6G (granulocytes, primarily neutrophils) to identify tumour-infiltrating leukocytes. Note that tertiary lymphoid structures, readily discernible adjacent to multiple lesions from both tumour-prone cohorts, were omitted from scoring to avoid skewing quantification. Whereas most tumour-infiltrating immune populations showed no quantitative difference between cohorts treated with or without asbestos, F4/80-positive macrophages were consistently and dramatically increased in tumours from asbestos-treated mice (Figure 4A, B).

**Figure 4.**
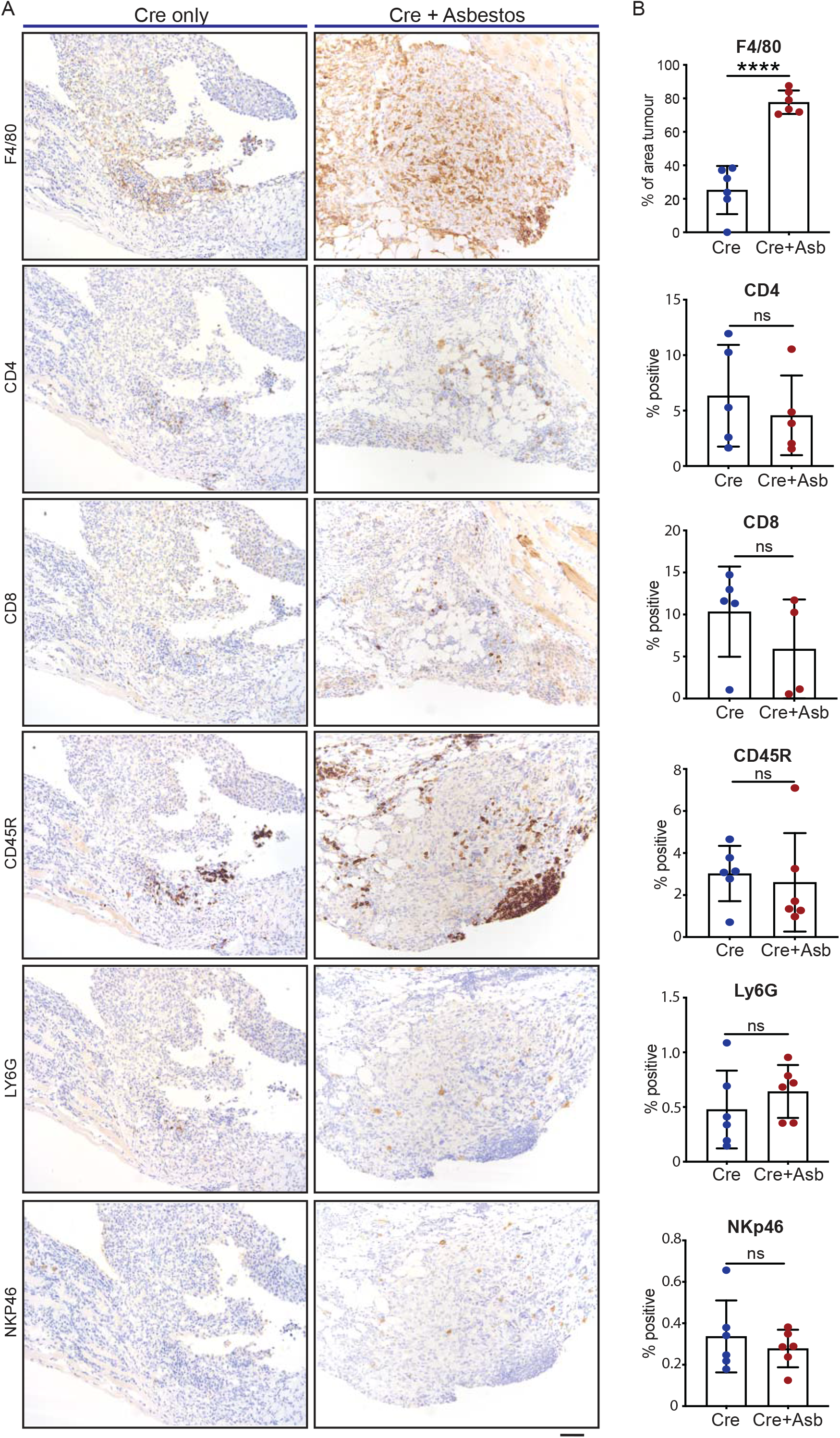
Asbestos drives increased macrophage infiltration in CNP tumours. **A)** Representative IHC images show detection of tumour infiltration by major leukocyte cell types in CNO mice induced with lenti-Cre with and without asbestos. Scale bar = 100μm. **B)** Quantification of tumour infiltrating leukocytes as per (A) in 6 mice from each cohort. **** denotes P < 0.0001 (T test); ns = not significant.

Similar to human MPM patients, pleural effusion was found in end-stage Cre-induced CNP mice and was more frequent in mice that received asbestos than those that did not. Pleural effusions are a rich source of inflammatory and effector immune populations and can harbour large numbers of viable tumour cells (27). Suspected cases of human MPM commonly first present with pleural effusion but only a relatively low percentage of these progress to MPM within 1-2 years (3). To investigate this disease feature further, we characterised immune cell infiltrates in the thoracic cavity of Cre-induced CNP mice treated with asbestos and harvested at 30 or 60 days post induction, ie. before the onset of any overt symptoms. At sacrifice, neoplastic lesions were visible in 0/6 mice harvested at 30 and 6/6 mice harvested at 60 days post induction. In order to normalise for variable effusion volume, we opted to perform a pleural lavage using 1ml of PBS. Recovered cell populations were stained with panels of myeloid and lymphoid marker antibodies and subject to flow cytometry for quantification. All major myeloid and lymphoid cell populations were readily detected in the pleural lavage from all CNP mice harvested at either 30 or 60 days post induction. Notably, the pleural lavage from 3 out of 6 mice harvested at d60 revealed a pronounced increase in the absolute numbers of all major leukocyte populations examined (Fig S2A-C). When expressed as a proportion of all infiltrating leukocytes, we note that macrophages and B cells show a pronounced increase from 30 to 60 days post induction, whereas eosinophils show a relative decrease within the same time frame (Fig S2D).

### Single-agent suppression of macrophages fails to impact survival

The CSF1 receptor regulates multiple aspects of macrophage function as well as survival, proliferation and differentiation from monocyte precursors (28). Given the pronounced increase in tumour infiltrating F4/80-positive macrophages in asbestos treated CNP mice, we asked if macrophage suppression could alone impact overall survival of this cohort. Starting from day 60 post induction, mice were treated twice daily with the selective CSF1R inhibitor AZD7507 (29), or vehicle control, and maintained on drug until symptoms required euthanasia (Fig. 5A). IHC for F4/80 confirmed suppression of tumour infiltration by macrophages in all mice treated with AZD7507, however no survival benefit was observed (Fig. 5B, C). We conclude that although asbestos exposure correlates with a sharp increase in tumour-infiltrating macrophages, single-agent suppression of macrophages by CSF1R blockade is ineffective for treatment of mesothelioma.

**Figure 5.**
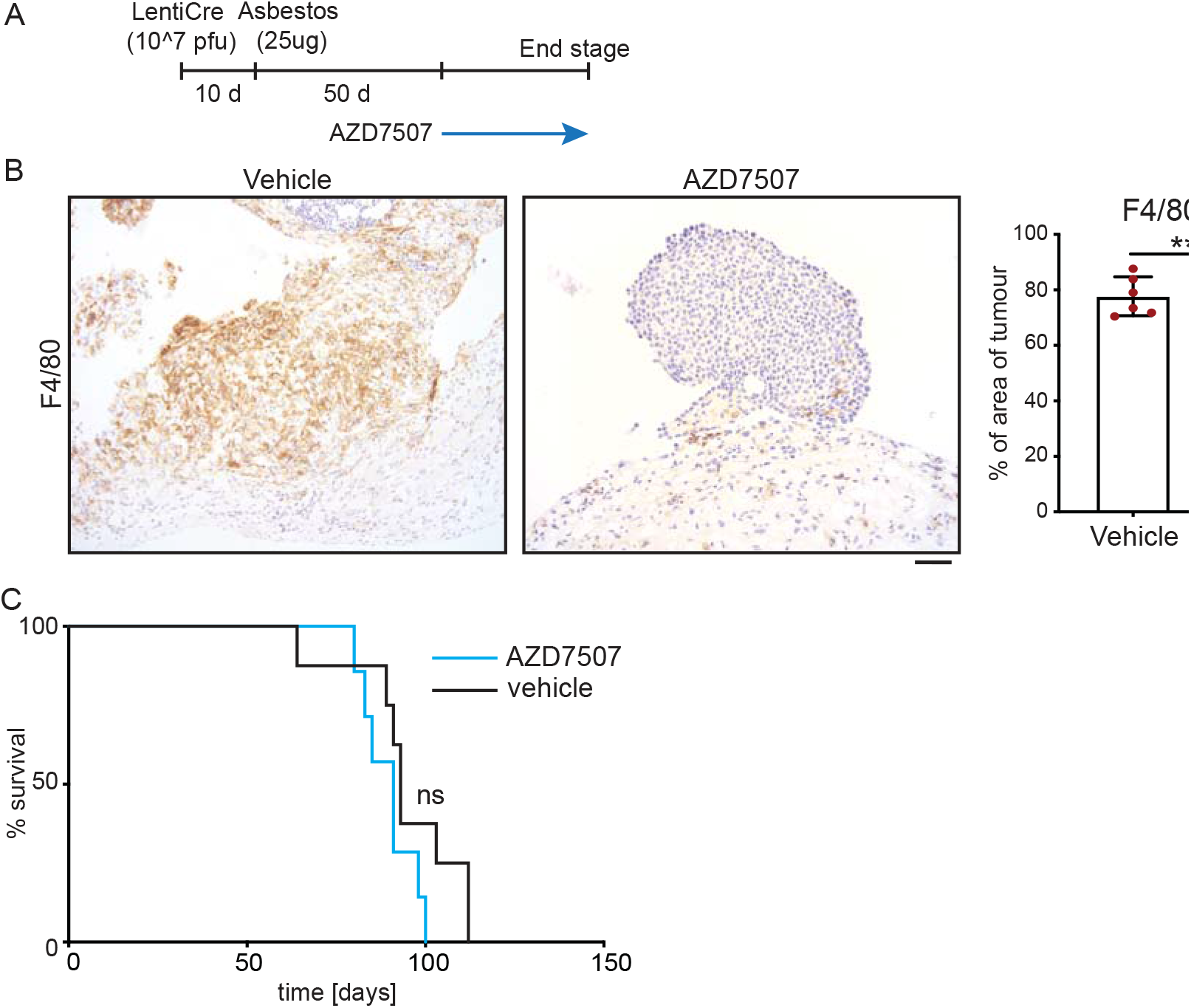
Single agent suppression of macrophages fails to impact survival of CNP mice. **A)** Schematic of intervention protocol. **B)** IHC validation of macrophage suppression by AZD7507. Left panels show representative images of F4/80-stained diaphragm tumours from end-stage CNP mice induced with lenti-Cre + asbestos and treated with AZD7507 (right) or vehicle control (left). Scale bar = 100μm. Right panel shows quantification of diaphragm tumour F4/80 staining in 5 mice from each cohort. **C)** Kaplan Meier plot of overall survival of mice treated with AZD7507 (N=7) or vehicle control (N=8). Ns = Not significant (Mantel Cox logrank test).

## DISCUSSION

Chronic inflammation plays an irrefutable, albeit poorly understood, role in the development of malignant pleural mesothelioma. The persistence of durable high-aspect ratio asbestos fibres sustains the chronically inflamed state throughout the patient history, from exposure all the way through presentation, treatment, and ultimately progression to lethal disease (9). However, the demonstration, in complex genetically engineered mice, that targeted deletion of combinations of tumour suppressor genes commonly lost in human MPM suffices to drive mesothelioma in the absence of asbestos exposure, indicates that chronic inflammation is not required *per se* for MPM development. A plausible conjecture is thus that chronic pleural inflammation in humans creates a permissive environment for such mutations to arise but plays no further role in disease progression. Here we directly investigated this assumption using the same combination of alleles previously shown to suffice for mesothelioma development in mice (19). We verified that Cre-induced CNP mice are genetically proficient for development of MPM in the absence of asbestos exposure. However, we found that the additional insult of asbestos-driven chronic inflammation dramatically accelerates the onset of lethal disease. While we cannot exclude the possibility that this acceleration derives from additional asbestos-driven mutations, we note that only 4 of 15 triple heterozygous CNP mice succumbed to mesothelioma within 15 months of induction with the combination of lenti-Cre + asbestos, which is consistent with the reported incidence for mesothelioma development in asbestos injected wild-type mice (7): If asbestos were driving widespread genomic instability, one would have expected loss of heterozygosity at one of more of these haplo-insufficient tumour suppressor genes to manifest as increased/accelerated incidence of malignancy.

At the cellular level, the inclusion of asbestos did not measurably influence the tumour phenotype, although differences in the apparent growth pattern of individual lesions were noted. We detected no consistent difference in cell proliferation, expression of epithelial or fibroblast lineage markers, nor indeed in the expression of WT1, in size-matched tumours from Cre-induced CNP mice with or without exposure to asbestos. In contrast, asbestos exposure was associated with dramatically increased presence of F4/80-positive macrophages in tumour masses. Macrophages are known to be early responders to asbestos, infiltrating into the chest cavity of mice within 48hrs of asbestos inoculation (8). Their inability to clear high aspect-ratio fibres is thought to establish the chronically inflamed state (7, 9). Macrophages can have either tumour suppressive or tumour-promoting activities (30), play major roles in recruitment or indeed suppression of other immune cell types (31), and are associated with resistance to radio- and chemotherapy as well as to certain targeted therapies (32). Although macrophage recruitment has been reported in GE mouse models of MPM without asbestos exposure (21) our data suggest that tumour-infiltrating macrophages are significantly under-represented in such models. Targeted suppression of macrophages failed to improve survival in our mouse model, consistent with a similar failure as single-agent treatment in other disease models and appropriately reflecting the challenge of effective treatment of this cancer. It remains to be determined if macrophage suppression can be used to enhance other treatment modalities, as has been shown for other cancer types (32).

The mouse model of mesothelioma presented here represents a significant refinement over previous models in 2 important ways. Firstly, our allele induction protocol did not give rise to off-target tumours, such as those reported using similar (20, 21) or indeed identical (19) allelic mice induced with high-dose adenoviral-Cre vectors, which include sarcomas, lymphomas, and, in the case of *Bap1* floxed mice, hepatocellular carcinomas. It is unclear if this difference arises from our use of a lentiviral rather than an adenoviral vector, the lower dose of virus we administered (10^7^ pfu Lenti-Cre compared with 10^9^ pfu Adeno-Cre), or a combination of both. The integration of lentiviral vectors into host cell genomes facilitates persistent expression of Cre recombinase, increasing the efficiency of floxed allele targeting and reducing the dose of virus required for effective Cre delivery. The caveat to such integration is of course that it can disrupt endogenous gene expression at the integration site. On the other hand, adenoviral vectors are inherently more stable than lentiviruses: While the tropism of lenti and adenoviral vectors would be broadly similar, the protein capsid of adenoviral vectors renders them considerably more robust than lentiviral vectors, and thus more likely to persist in circulation long enough to infect unintended cell populations (33).

The second major distinction is the epithelioid histology of tumours arising in our mouse model, irrespective of asbestos exposure, as compared with the bi-phasic and sarcomatoid histologies of previously reported models, including those using the same allelic combination. As is the case for off-target tumour incidence, we suspect that choice and/or dose of vector may largely explain the difference in tumour histology. Specifically, adenoviral vectors are profoundly more immunogenic than lentiviral vectors and this immunogenicity may well influence the trajectory of the nascent tumour phenotype (33). Notably, whereas loss of *BAP1* is enriched in human epithelioid MPM (13), the recently reported adeno-Cre induced mouse models incorporating a floxed *Bap1* allele both yielded sarcomatoid phenotypes (20, 21). Conversely, loss of *NF2* and *Tp53* are associated with the sarcomatoid subtype in human MPM (12), yet our model, which incorporates both of these tumour suppressors, yielded epithelioid tumours. These observations lead us to speculate that the microenvironment present during tumour initiation may have a stronger influence over tumour histology than the presence of any given mutation. We note that a similar phenomenon of differential tumour phenotypes arising in genetically identical models of liver cancer was recently reported: activation of the same oncogenic drivers yielded either hepatocellular carcinoma or intrahepatic cholangiocarcinoma, depending on whether cells neighbouring the tumour stem cells died via apoptosis or via necroptosis, respectively, during allele induction (34). It is well-established that necroptosis is a much more inflammatory form of cell death than apoptosis (35) – these insights likely have direct relevance both for influencing how mesothelioma subtypes are modelled in mice and potentially for how the disease manifests in humans.

It is also clear that oncogenic mutations profoundly shape the tumour microenvironment and that different mutations will have distinct impacts on tumour-cell expression of cytokines, chemokines and other factors that regulate these effects (13, 36, 37). Our model now provides a new platform for investigating how recurring mutations in mesothelioma influence the disease-relevant inflammatory response to asbestos and *vice versa*. The inclusion of asbestos additionally addresses a vital aspect of immediate relevance for modelling the human disease in GE mice through more accurate recapitulation of the inflammatory microenvironment that is such a crucial part of this disease. Giving particular consideration to the emerging prominence of immunotherapy in mesothelioma (38), we believe that this model will provide a more accurate platform for pre-clinical evaluation of new therapeutic agents and strategies.

## MATERIALS AND METHODS

### Mice & Procedures

The *Nf2* floxed (39), *Tp53* floxed (40) and *Cdkn2a* conditional knockout (41) mice were described previously. All experiments involving mice were approved by the local ethics committee and conducted in accordance with UK Home Office license numbers PE47BC0BF, 70/7950 & 70/8646. Experiments were performed and reported in accordance with the ARRIVE guidelines. Mice were housed in a constant 12hr light/dark cycle and fed/watered *ad libitum*. Both males and females were included in approximately equal numbers and mice were randomly assigned to induction or treatment cohorts, balanced only for sex. Recombinant lentivirus expressing Cre recombinase was purchased from the University of Iowa vector core facility. To induce allele recombination, administration of lentiviral vector carrying CRE recombinase was performed on mice aged 8-10 weeks via single intrapleural injection of 10^7^ viral particles per mouse. Where indicated, asbestos (long fibre amosite) was administered via single intrapleural injection of 25μg fibres per mouse 10 days post CRE administration. CSF1R inhibitor (AZ7507) was provided by the manufacturer under a research collaboration agreement with CRUK Beatson Institute. AZ7507 was dissolved in 0.5% HPMC + 0.1% Tween80 in dH_2_O and administered at 100mg/kg twice daily by oral gavage. Routine health monitoring was performed by facility personnel without knowledge of experimental details. Humane end points were defined as exhibition of 2 or more symptoms: weight loss, elevated breathing, hunching, untidy coat. All mice were sacrificed using a schedule 1 procedure.

### Histology & Immunohistochemistry

For histological examination, tissue was harvested, fixed overnight in formalin and embedded in paraffin for sectioning, followed by standard staining in hematoxylin and eosin (H&E). Tumour pathology was determined by an experienced translational thoracic pathologist (JLQ). For manual IHC staining for Ki67 (Fisher Scientific, RM-9106-S0, 1:1000), FFPE tissue sections were deparaffinized in 3 changes of xylene and rehydrated in graded ethanol solutions. Antigen retrieval was performed by microwaving in 10mM Sodium Citrate, pH6.0. Endogenous peroxidases were quenched in 3% H_2_O_2_ and non-specific binding was blocked with 1 or 3% BSA solution. Staining for all other antibodies (see table below) were performed by Leica Bond Rx or Dako autostainers: Sections were dewaxed and antigen retrieval was performed on board, ER2 for 20 mins at 95°C or Enz1 for 10 mins at 37°C. For Dako autostaining, pH6 antigen retrieval was performed off-board in a Dako PT module using the pH6 retrieval buffer for 20 mins at 97°C. High pH TRS (pH9) was done off-board in a Dako PT module using the high pH retrieval buffer for 20 mins at 97C. Liquid DAB was used as chromogen in all cases. Detailed antibody information and staining conditions are as follows:

| Ab | Clone | Company | Code | Autostainer | Retrieval | Dilution |
| --- | --- | --- | --- | --- | --- | --- |
| B220/CD45R | RA3-6B2 | Abcam | Ab64100 | Leica Bond Rx | ER2 20 mins | 1:200 |
| CD4 | 4SM 95 | eBioscience | 14-9766-82 | Leica Bond Rx | ER2 20 mins | 1:500 |
| CD8 | 4SM15 | eBioscience | 14-0808-82 | Leica Bond Rx | ER2 20 mins | 1:500 |
| Cytokeratin (Pan) | AE1/AE3 | Thermo | MS-343 | Dako Autostainer Link48 | pH6 20 mins | 1:100 |
| F4/80 | CI:A3-1 | Abcam | Ab6640 | Leica Bond RX | Enz1 10 min | 1:100 |
| LY6G | IA8 | BioXcell | BE0075-1 | Leica Bond RX | ER2 20 mins | 1:60,000 |
| NKP46/NCR1 |  | R &D systems | af2225 | Dako Autostainer Link48 | High pH TRS | 1:200 |
| αSMA | 1A4 | Sigma-Aldrich | A2547 | Dako Autostainer Link48 | pH6 20 mins | 1:25000 |
| Vimentin | D21H3 | Cell Signalling | 5741 | Dako Autostainer Link48 | High pH TRS | 1:100 |
| WT1 | [CAN-R9(IHC)-56-2] | Abcam | Ab89901 | Dako Autostainer Link48 | High pH TRS | 1:250 |

### IHC Quantification

Quantification was performed on IHC stained slides using QuPath open source digital pathology software (42). Tumour regions were annotated manually followed by automated cell detection analysis. For F4/80 staining, accurate cell counts could not be obtained using QuPath and data are presented as area of tumour stained positively. Graphical representation of data was performed using GraphPad-Prism.

### Statistical analysis

Quantitative data were uploaded into Prism spreadsheets for analysis and graphic production. Statistical significance was determined by unpaired *t* test (ns = not significant; p>0.05, *; p<0.05, **; p<0.01, ***, p<0.001; ****, p<0.0001). For Kaplan-Meier plots, the Mantel Cox logrank test was performed.

## ACKNOWLEDGMENTS

We wish to thank Prof. Anton Berns (NKI) for generously providing the CNP mice used in this study and Seth Coffelt (University of Glasgow & CRUK Glasgow Cancer Centre) for extensive discussions and scientific insights. Special thanks to all the staff at the CRUK Beatson Biological Services Unit and Histology core facility.

## AUTHOR CONTRIBUTIONS

Project conception and study design: DJM, MMF, KB, JLQ

Performed experiments: PF, KG, CR, XRI, EJ

Data analysis & figure production: PF, KG, CR, XRI, EJ, LMC, DL, JLQ, MMF, DJM

Reagent & technique development: TC, TM, RD

Secured funding: DJM, KB, MMF, JLQ, CR

Text preparation and editing: All authors

## REFERENCES

1. Andujar P, Lacourt A, Brochard P, Pairon JC, Jaurand MC, Jean D. Five years update on relationships between malignant pleural mesothelioma and exposure to asbestos and other elongated mineral particles. J Toxicol Environ Health B Crit Rev. 2016;19(5–6):151–72.

2. Carbone M, Kanodia S, Chao A, Miller A, Wali A, Weissman D, et al. Consensus Report of the 2015 Weinman International Conference on Mesothelioma. J Thorac Oncol. 2016;11(8):1246–62.

3. Bibby AC, Tsim S, Kanellakis N, Ball H, Talbot DC, Blyth KG, et al. Malignant pleural mesothelioma: an update on investigation, diagnosis and treatment. Eur Respir Rev. 2016;25(142):472–86.

4. Tsao AS, Lindwasser OW, Adjei AA, Adusumilli PS, Beyers ML, Blumenthal GM, et al. Current and Future Management of Malignant Mesothelioma: A Consensus Report from the National Cancer Institute Thoracic Malignancy Steering Committee, International Association for the Study of Lung Cancer, and Mesothelioma Applied Research Foundation. J Thorac Oncol. 2018;13(11):1655–67.

5. Blyth KG, Murphy DJ. Progress and challenges in Mesothelioma: From bench to bedside. Respir Med. 2018;134:31–41.

6. Port J, Murphy DJ. Mesothelioma: Identical Routes to Malignancy from Asbestos and Carbon Nanotubes. Curr Biol. 2017;27(21):R1173–R6.

7. Chernova T, Murphy FA, Galavotti S, Sun XM, Powley IR, Grosso S, et al. Long-Fiber Carbon Nanotubes Replicate Asbestos-Induced Mesothelioma with Disruption of the Tumor Suppressor Gene Cdkn2a (Ink4a/Arf). Curr Biol. 2017;27(21):3302–14 e6.

8. Donaldson K, Brown GM, Brown DM, Bolton RE, Davis JM. Inflammation generating potential of long and short fibre amosite asbestos samples. Br J Ind Med. 1989;46(4):271–6.

9. Donaldson K, Murphy FA, Duffin R, Poland CA. Asbestos, carbon nanotubes and the pleural mesothelium: a review of the hypothesis regarding the role of long fibre retention in the parietal pleura, inflammation and mesothelioma. Part Fibre Toxicol. 2010;7:5.

10. Murphy FA, Schinwald A, Poland CA, Donaldson K. The mechanism of pleural inflammation by long carbon nanotubes: interaction of long fibres with macrophages stimulates them to amplify pro-inflammatory responses in mesothelial cells. Part Fibre Toxicol. 2012;9:8.

11. Benedetti S, Nuvoli B, Catalani S, Galati R. Reactive oxygen species a double-edged sword for mesothelioma. Oncotarget. 2015;6(19):16848–65.

12. Bueno R, Stawiski EW, Goldstein LD, Durinck S, De Rienzo A, Modrusan Z, et al. Comprehensive genomic analysis of malignant pleural mesothelioma identifies recurrent mutations, gene fusions and splicing alterations. Nat Genet. 2016;48(4):407–16.

13. Hmeljak J, Sanchez-Vega F, Hoadley KA, Shih J, Stewart C, Heiman D, et al. Integrative Molecular Characterization of Malignant Pleural Mesothelioma. Cancer Discov. 2018;8(12):1548–65.

14. Tallet A, Nault JC, Renier A, Hysi I, Galateau-Salle F, Cazes A, et al. Overexpression and promoter mutation of the TERT gene in malignant pleural mesothelioma. Oncogene. 2014;33(28):3748–52.

15. Marsella JM, Liu BL, Vaslet CA, Kane AB. Susceptibility of p53-deficient mice to induction of mesothelioma by crocidolite asbestos fibers. Environ Health Perspect. 1997;105 Suppl 5:1069–72.

16. Fleury-Feith J, Lecomte C, Renier A, Matrat M, Kheuang L, Abramowski V, et al. Hemizygosity of Nf2 is associated with increased susceptibility to asbestos-induced peritoneal tumours. Oncogene. 2003;22(24):3799–805.

17. Altomare DA, Menges CW, Xu J, Pei J, Zhang L, Tadevosyan A, et al. Losses of both products of the Cdkn2a/Arf locus contribute to asbestos-induced mesothelioma development and cooperate to accelerate tumorigenesis. PLoS One. 2011;6(4):e18828.

18. Kadariya Y, Cheung M, Xu J, Pei J, Sementino E, Menges CW, et al. Bap1 Is a Bona Fide Tumor Suppressor: Genetic Evidence from Mouse Models Carrying Heterozygous Germline Bap1 Mutations. Cancer Res. 2016;76(9):2836–44.

19. Jongsma J, van Montfort E, Vooijs M, Zevenhoven J, Krimpenfort P, van der Valk M, et al. A conditional mouse model for malignant mesothelioma. Cancer Cell. 2008;13(3):261–71.

20. Kukuyan AM, Sementino E, Kadariya Y, Menges CW, Cheung M, Tan Y, et al. Inactivation of Bap1 Cooperates with Losses of Nf2 and Cdkn2a to Drive the Development of Pleural Malignant Mesothelioma in Conditional Mouse Models. Cancer Res. 2019;79(16):4113–23.

21. Badhai J, Pandey GK, Song JY, Krijgsman O, Bhaskaran R, Chandrasekaran G, et al. Combined deletion of Bap1, Nf2, and Cdkn2ab causes rapid onset of malignant mesothelioma in mice. J Exp Med. 2020;217(6).

22. Rossini M, Rizzo P, Bononi I, Clementz A, Ferrari R, Martini F, et al. New Perspectives on Diagnosis and Therapy of Malignant Pleural Mesothelioma. Front Oncol. 2018;8:91.

23. Lee HS, Jang HJ, Choi JM, Zhang J, de Rosen VL, Wheeler TM, et al. Comprehensive immunoproteogenomic analyses of malignant pleural mesothelioma. JCI Insight. 2018;3(7).

24. Patil NS, Righi L, Koeppen H, Zou W, Izzo S, Grosso F, et al. Molecular and Histopathological Characterization of the Tumor Immune Microenvironment in Advanced Stage of Malignant Pleural Mesothelioma. J Thorac Oncol. 2018;13(1):124–33.

25. Klampatsa A, O’Brien SM, Thompson JC, Rao AS, Stadanlick JE, Martinez MC, et al. Phenotypic and functional analysis of malignant mesothelioma tumor-infiltrating lymphocytes. Oncoimmunology. 2019;8(9):e1638211.

26. Blum Y, Meiller C, Quetel L, Elarouci N, Ayadi M, Tashtanbaeva D, et al. Dissecting heterogeneity in malignant pleural mesothelioma through histo-molecular gradients for clinical applications. Nat Commun. 2019;10(1):1333.

27. Minnema-Luiting J, Vroman H, Aerts J, Cornelissen R. Heterogeneity in Immune Cell Content in Malignant Pleural Mesothelioma. Int J Mol Sci. 2018;19(4).

28. Chitu V, Gokhan S, Gulinello M, Branch CA, Patil M, Basu R, et al. Phenotypic characterization of a Csf1r haploinsufficient mouse model of adult-onset leukodystrophy with axonal spheroids and pigmented glia (ALSP). Neurobiol Dis. 2015;74:219–28.

29. Candido JB, Morton JP, Bailey P, Campbell AD, Karim SA, Jamieson T, et al. CSF1R(+) Macrophages Sustain Pancreatic Tumor Growth through T Cell Suppression and Maintenance of Key Gene Programs that Define the Squamous Subtype. Cell Rep. 2018;23(5):1448–60.

30. Zhang QW, Liu L, Gong CY, Shi HS, Zeng YH, Wang XZ, et al. Prognostic significance of tumor-associated macrophages in solid tumor: a meta-analysis of the literature. PLoS One. 2012;7(12):e50946.

31. Biswas SK, Mantovani A. Macrophage plasticity and interaction with lymphocyte subsets: cancer as a paradigm. Nat Immunol. 2010;11(10):889–96.

32. Ruffell B, Coussens LM. Macrophages and therapeutic resistance in cancer. Cancer Cell. 2015;27(4):462–72.

33. Lundstrom K. Viral Vectors in Gene Therapy. Diseases. 2018;6(2).

34. Seehawer M, Heinzmann F, D’Artista L, Harbig J, Roux PF, Hoenicke L, et al. Necroptosis microenvironment directs lineage commitment in liver cancer. Nature. 2018;562(7725):69–75.

35. Vringer E, Tait SWG. Mitochondria and Inflammation: Cell Death Heats Up. Front Cell Dev Biol. 2019;7:100.

36. Chiasson-MacKenzie C, Morris ZS, Liu CH, Bradford WB, Koorman T, McClatchey AI. Merlin/ERM proteins regulate growth factor-induced macropinocytosis and receptor recycling by organizing the plasma membrane:cytoskeleton interface. Genes Dev. 2018;32(17-18):1201–14.

37. Muthalagu N, Monteverde T, Raffo-Iraolagoitia X, Wiesheu R, Whyte D, Hedley A, et al. Repression of the Type I Interferon Pathway Underlies MYC- and KRAS-Dependent Evasion of NK and B Cells in Pancreatic Ductal Adenocarcinoma. Cancer Discov. 2020;10(6):872–87.

38. Scherpereel A, Mazieres J, Greillier L, Lantuejoul S, Do P, Bylicki O, et al. Nivolumab or nivolumab plus ipilimumab in patients with relapsed malignant pleural mesothelioma (IFCT-1501 MAPS2): a multicentre, open-label, randomised, non-comparative, phase 2 trial. Lancet Oncol. 2019;20(2):239–53.

39. Giovannini M, Robanus-Maandag E, van der Valk M, Niwa-Kawakita M, Abramowski V, Goutebroze L, et al. Conditional biallelic Nf2 mutation in the mouse promotes manifestations of human neurofibromatosis type 2. Genes Dev. 2000;14(13):1617–30.

40. Marino S, Vooijs M, van Der Gulden H, Jonkers J, Berns A. Induction of medulloblastomas in p53-null mutant mice by somatic inactivation of Rb in the external granular layer cells of the cerebellum. Genes Dev. 2000;14(8):994–1004.

41. Krimpenfort P, Quon KC, Mooi WJ, Loonstra A, Berns A. Loss of p16Ink4a confers susceptibility to metastatic melanoma in mice. Nature. 2001;413(6851):83–6.

42. Bankhead P, Loughrey MB, Fernandez JA, Dombrowski Y, McArt DG, Dunne PD, et al. QuPath: Open source software for digital pathology image analysis. Sci Rep. 2017;7(1):16878.

