## Supplemental figures and methods for "Asbestos accelerates disease onset in a genetic model of Malignant Pleural Mesothelioma"

**List of Supplemental Material**

Figure S1. FDG uptake by mesothelioma in CNP mice induced with lenti-Cre + asbestos

Figure S2. Detection of major leukocyte populations in pleural lavage from pre-symptomatic CNP mice induced with lenti-Cre + asbestos

Supplemental Methods

Figure S1

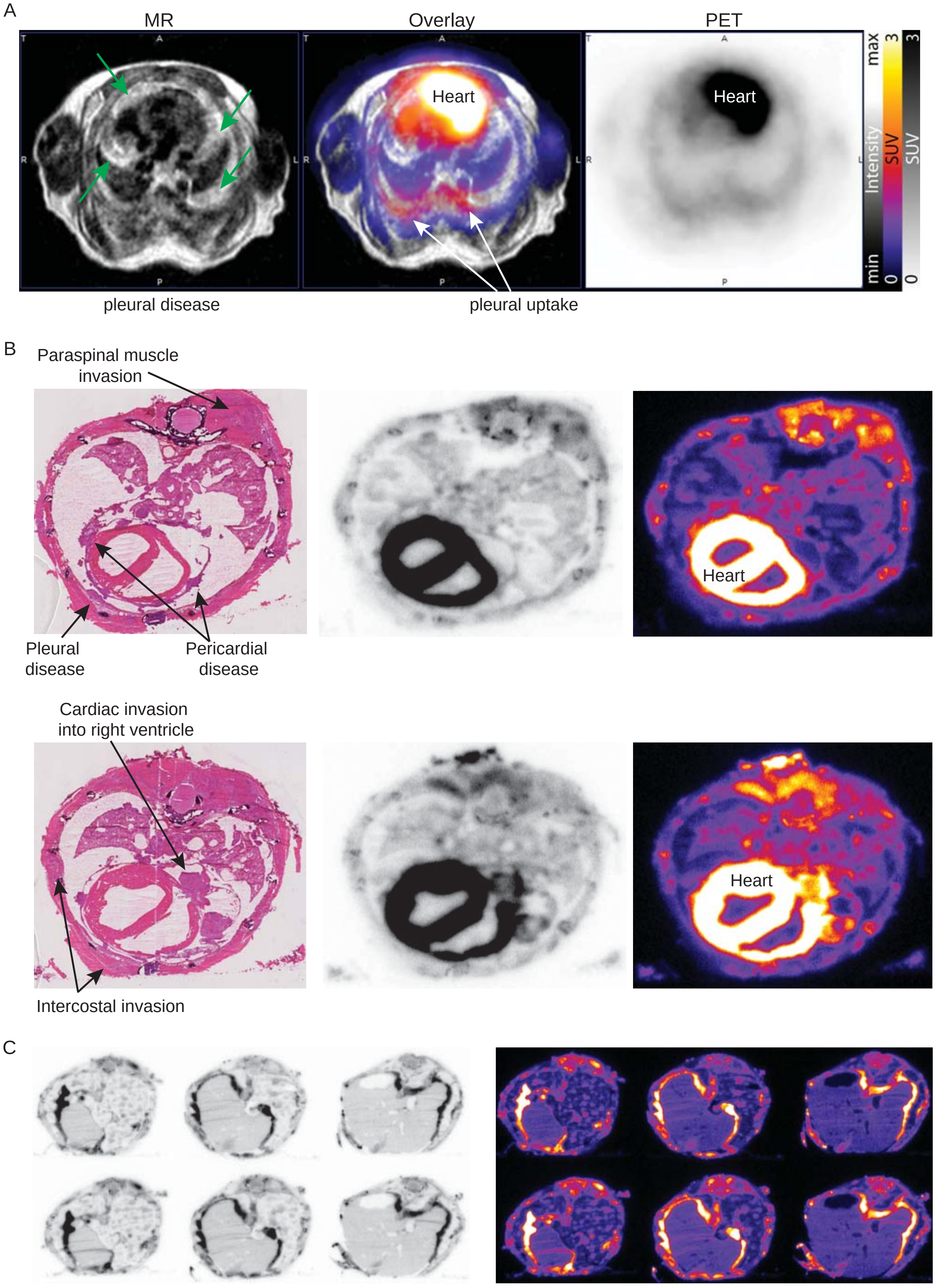
A
MR
Overlay
PET
Heart
Heart
max
min
Intensity
SUV
3
0
SUV
3
0
pleural disease
pleural uptake
B
Paraspinal muscle invasion
Pleural disease
Pericardial disease
Heart
Cardiac invasion into right ventricle
Heart
Intercostal invasion
C

Figure S2

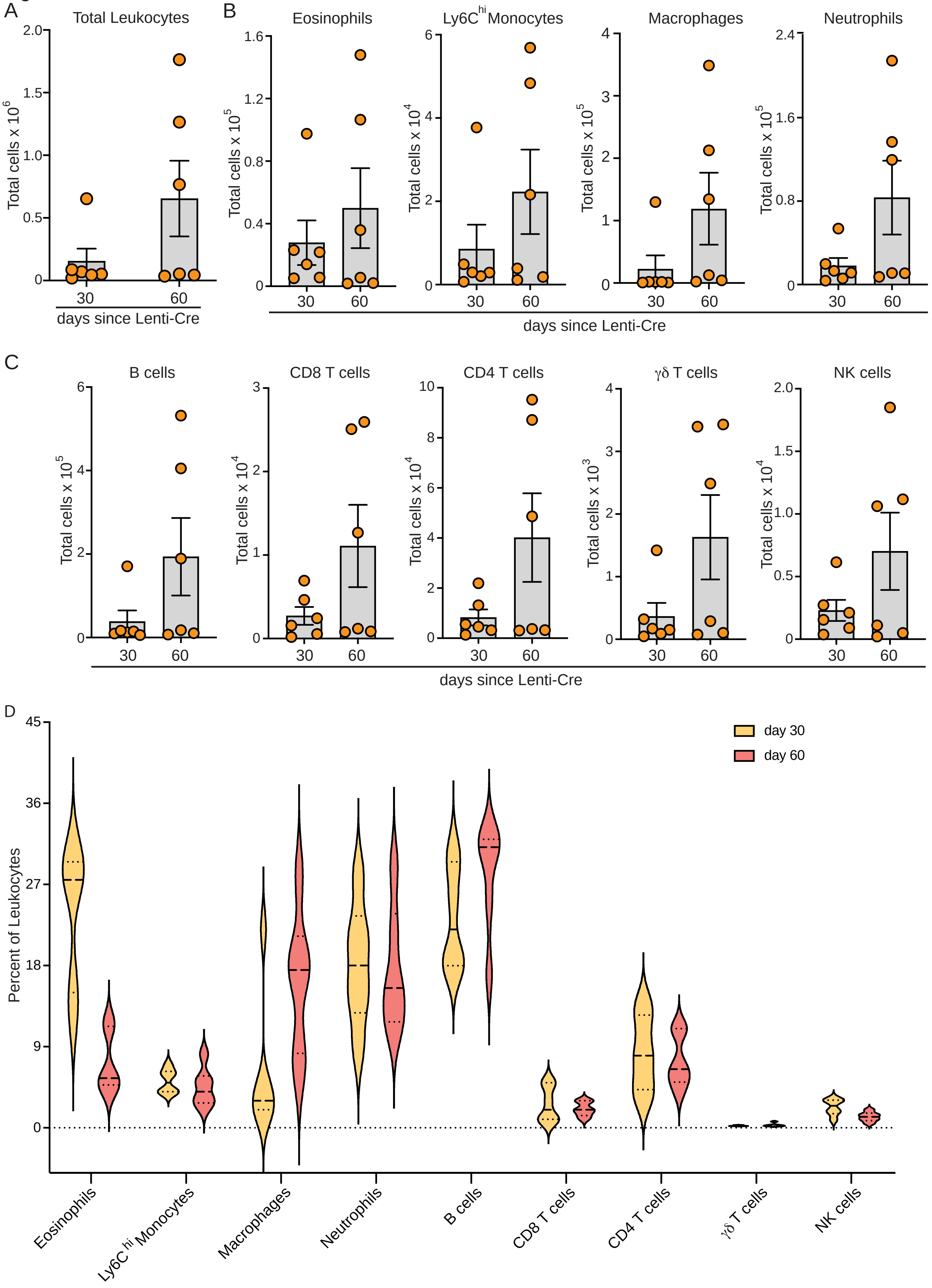

A
Total Leukocytes
Total cells x 10^6
2.0
1.5
1.0
0.5
0
30
60
days since Lenti-Cre
B
Eosinophils
Ly6C hi Monocytes
Macrophages
Neutrophils
Total cells x 10^5
Total cells x 10^4
1.6
1.2
0.8
0.4
6
4
2
3
2.4
30
60
days since Lenti-Cre
C
B cells
CD8 T cells
CD4 T cells
γδ T cells
NK cells
Total cells x 10^3
10
8
days since Lenti-Cre
D
day 30
day 60
Percent of Leukocytes
45
36
27
18
9
Eosinophils
Ly6C hi Monocytes
Macrophages
Neutrophils
B cells
CD8 T cells
CD4 T cells
γδ T cells
NK cells

**SUPPLEMENTAL FIGURE LEGENDS**

**Figure S1. FDG uptake by mesothelioma in CNP mice induced with lenti-Cre + asbestos**

**A)** FDG PET/MR imaging of CNP mice induced with lenti-Cre + asbestos at end-stage. Left: T2 FSE 3D axial MR of thoracic cavity. Green arrows indicate abnormal pleural thickening. Middle: T2 FSE 3D axial MR co-registered with false-colour PET image showing uptake of [$^{18}$F]-FDG in the pleura. Right: grey-scale [$^{18}$F]-FDG PET image. **B)** Transverse sections through the chest cavity showing H&E stained tissue (left panels) and [$^{18}$F]-FDG detection by autoradiography in brightfield (centre panels) and false-colour (right panels) of corresponding whole mount sections. Invasive, pericardial and pleural disease indicated by arrows. **C)** Autoradiography of [$^{18}$F]-FDG detection showing extent of pleural disease in an exemplar mouse in brightfield (left) and false colour (right). Note detection of pleural effusion in the 3$^{rd}$ section from left (white area in brightfield, black area in false colour).

**Figure S2. Detection of major leukocyte populations in pleural lavage from pre-symptomatic CNP mice induced with lenti-Cre + asbestos**

**A)** Total CD45$^{+}$ WBC counts from CNP mice harvested at 30 days (N=6) and 60 days (N=6) post lenti-Cre induction, measured by FACS analysis. **B)** Major myeloid lineage populations sorted from (A). **C)** Major lymphoid lineage populations sorted from (A). **D)** Major WBC populations expressed as a percentage of total from (A).

**SUPPLEMENTAL METHODS**

**PET/MR Imaging**

PET/MR imaging was based on a published protocol (42). In brief, we anaesthetised and maintained mice using 1-3% isoflurane in 2L/min oxygen before cannulating the lateral tail vein. Mouse body temperature was maintained at 37$^{o}$C using an external heat source prior to and throughout anaesthesia and imaging. A bolus intravenous injection of 200ul [$^{18}$F]FDG (41.71±9.9MBq), diluted in 0.9% saline solution, was administered via the tail vein cannula. We acquired proton T2-weighted MR images (T2 FSE 3D axial MR, resolution 0.13x0.13x0.5, matrix size 168x160x150, Repetition time (TR) 2000msec, Echo time (TE) 54.6msec, Flip Angle 90 degrees, Number of excitations 2), in the prone position during an anaesthetised uptake phase of 50 mins, followed by a PET scan at 50-60mins using a NanoScan PET/MRI (1T) scanner (Mediso Ltd, Budapest, Hungary). PET images were corrected for decay, attenuation, scatter, deadtime and random coincidences and reconstructed (3D iterative static reconstruction, matrix size 105x105x237, with isotropic 0.4mm voxels) using 3D Tera-Tomo (Mediso Ltd, Budapest, Hungary). Images were visualised using Vivoquant version 3.0 (InviCRO, MA, USA).

Immediately following PET, mice were euthanized by $CO_2$ inhalation. A small incision was made in the trachea and a blunt 19G cannula (Western laboratory service Ltd, Horndean, UK) was inserted. A 90% OCT/10% PBS mix was drawn into a 5ml syringe which was then slowly dispensed into the lungs. Once the lungs were fully inflated, the cannula was removed and the jugular vein was cut to allow for blood expansion during freezing. The carcass was then placed in 2-methybutane (Sigma-Aldrich, MO, USA) which had been cooled for 5 mins on dry ice. Once frozen, the fur was removed and the torso was isolated and mounted on a specimen disc, using Optimal Cutting Temperature (OCT) solution (Tissue-Tek, CA, USA) for cutting on a Leica CM3050 S cryostat (Leica Biosystems, Wetzlar, Germany). Sections were cut at 10µm and mounted on PolyFrost Poly Lysine Coated Adhesive Frosted slides (Solmedia Ltd, Shrewsbury, UK) which were then placed face down on a phosphor screen BAS-IP SR 2 (GE Healthcare Lifescience, MA, USA), and left overnight in an exposure cassette (GE Healthcare Lifescience, MA, USA). The following day, phosphor screens were scanned using a Typhoon FLA 7000 IP imager (GE Healthcare Lifesciences, MA, USA) using a 635 nm excitation laser and an IP (390 BP) filter. Image J (National Institutes of Health, MD, USA) was used to visualise the autoradiography sections. We stained the sections with H&E, and scanned them using a Leica SCN400F slide scanner (Leica Biosystems, Wetzlar, Germany) and aligned these images to the autoradiograms acquired from the Typhoon.

**Flow cytometry**

Cells from pleural lavages were collected and split into two wells to be stained for flow cytometry as follows. After red blood cell lysis, cells were stained with ZombieNIR (Biolegend, 423106) and then blocked with TruStain FcX (Biolegend, 101320). Next, cells were incubated with fluorescently conjugated Abs (see table below) diluted in Brilliant Stain Buffer (BD, 566349) and fixed in 2% formaldehyde in PBS (Thermo, 28908). Before analysis using a BD Fortessa flow cytometer, counting beads (Spherotech, Catalog # ACFP-70-10) were added for quantification. Data were analysed with FlowJo software V.10. (BD). Leukocytes (Single cells ZombieNIR$^{-}$CD45$^{+}$) were further gated using panel I as eosinophils (CD11b$^{+}$SiglecF$^{+}$), Ly6C$^{hi}$ monocytes (SiglecF$^{-}$CD11b$^{+}$CD115$^{+}$F4/80$^{-}$Ly6C$^{hi}$), macrophages (SiglecF$^{-}$CD11b$^{+}$CD115$^{+}$F4/80$^{+}$) and neutrophils (SiglecF$^{-}$CD11b$^{+}$CD115$^{-}$Ly6G$^{+}$); and panel II as B cells (CD19$^{+}$CD3$^{-}$), gd T cells (CD19$^{-}$CD3$^{+}$TCRD$^{-}$), CD8 T cells (CD19$^{-}$CD3$^{+}$TCRD$^{-}$CD8$^{+}$), CD4 T cells (CD19$^{-}$CD3$^{+}$TCRD$^{-}$CD8$^{-}$CD4$^{+}$) and NK cells (CD19$^{-}$CD3$^{-}$TCRD$^{-}$CD8$^{-}$CD4$^{-}$NKp46$^{+}$). Antibodies used for FACS analysis listed in the table below:

| Panel | Ab | Clone | Company | Code |
|---|---|---|---|---|
| I | CD115-BV421 | AFS98 | Biolegend | 135513 |
| I | CD11b-BV650 | M1/70 | Biolegend | 101259 |
| II | CD19-BV605 | 6D5 | Biolegend | 115540 |
| II | CD3-Percp/Cy5.5 | 145-2C11 | Biolegend | 100328 |
| II | CD4-PE/CY7 | RM4-4 | Biolegend | 116016 |
| I-II | CD45-BV711 | 30-F11 | Biolegend | 103147 |
| II | CD8-BUV395 | 53-6.7 | BD Bioscience | 563786 |
| I | F4/80-FITC | Cl:A3-1 | BIO-RAD | MCA497F |
| I | Ly6C-Percp/Cy5.5 | HK1.4 | Biolegend | 128012 |
| I | Ly6G-BUV395 | 1A8 | BD Bioscience | 563978 |
| II | NKp46-BV421 | 29A1.4 | Biolegend | 137612 |
| I | SiglecF-AF647 | E50-2440 | BD Bioscience | 562680 |
| II | TCRD-FITC | GL3 | Biolegend | 118105 |
